# Stable, Easy-to-Handle, Fully Autologous Electrospun Polymer-Peptide Skin Equivalent for Severe Burn Injuries

**DOI:** 10.1101/2025.01.12.632550

**Authors:** Dana Cohen-Gerassi, Marina BenShoshan, Adi Liiani, Tomer Reuveni, Offir Loboda, Moti Haratz, Josef Haik, Itzhak Binderman, Yosi Shacham-Diamand, Amit Sitt, Ayelet Di Segni, Lihi Adler-Abramovich

**Affiliations:** Department of Oral Biology, The Goldschleger School of Dental Medicine, Faculty of Medical & Health Sciences, Tel Aviv University, Tel Aviv 6997801, Israel; Jan Koum Center for Nanoscience and Nanotechnology, Tel Aviv University, Tel Aviv 6997801, Israel; Department of Materials Science and Engineering, Tel Aviv University, Tel Aviv 6997801, Israel; Green Skin Engineering Laboratory, The Division of Plastic Surgery and the Intensive Care Burn Unit, Sheba Medical Center, Tel Hashomer, Israel; Department of Physical Chemistry, The School of Chemistry, Raymond and Beverly Sackler Faculty of Exact Sciences, Tel-Aviv University, Tel Aviv 6997801, Israel; The Division of Plastic & Reconstructive Surgery and The Intensive Care Burn Unit, Sheba Medical Center, Israel; The Scojen Institute for Synthetic Biology, Director, Reichman University, 8 University St., Herzliya 4610101, Israel

## Abstract

Severe burn injuries represent a significant clinical challenge due to their complex healing process and the high risk of complications, including infection, scarring, and contracture formation. Current therapeutic approaches for burn wound treatment include autologous donor-site grafting and advanced cell therapy techniques like cultured epidermal autografts (CEA), which successfully facilitate wound closure through re-epithelialization. However, CEAs are limited by fragility, shrinkage, lack of a dermal layer, and risks of contamination. Here, aiming to overcome these limitations, we developed a personalized skin equivalent featuring an engineered scaffold composed of electrospun polycaprolactone (PCL) functionalized with the bioactive peptide fluorenylmethyloxycarbonyl-phenylalanine-arginine-glycine-aspartic acid (Fmoc-FRGD). This scaffold is designed to mimic the natural extracellular matrix (ECM), promoting cellular adhesion, integration, and proliferation while maintaining structural integrity. *In-vitro* analysis demonstrated the scaffold’s ability to support multi-layered human skin cell growth, while *in-vivo* experiments confirmed its efficacy in facilitating wound closure and full-thickness skin regeneration in a murine model. This bioengineered skin equivalent is mechanically robust, easy to handle, fully autologous and exhibits no contraction, offering a transformative therapeutic alternative for the treatment of severe burn injuries.

## Introduction

Severe burn wounds present a major global health issue, affecting millions annually and resulting in high morbidity, scarring, limited mobility, and prolonged recovery.^[1]^ These injuries are characterized by the loss of skin structure, function, and regenerative capacity. Despite advancements in medical treatments, effective solutions for restoring skin function and regeneration in extensive burn cases remain limited.^[2]^

For nearly four decades, the treatment for extensive burn wounds has relied heavily on donor autologous skin grafts, and cultured epidermal autografts (CEA). While CEA facilitates re-epithelialization and improves wound closure, it has several limitations including,^[3,4]^ (1) The mechanical fragility of CEA sheets is a notable concern, making them difficult to handle and apply. Furthermore, the absence of a dermal layer compromises structural support, resulting in lower graft take rates and increasing the risk of functional deficits and poor aesthetic outcomes.^[5]^ (2) CEAs require a long preparation time and high production costs; it takes 2–4 weeks to culture the sheets under strict sterile conditions to prevent contamination and ensure successful cell growth.^[6]^ (3) The final grafts may contract by more than half of their original size upon detachment from the culture dish.^[7]^ (4) CEAs require a feeder layer for its culturing. This feeder layer is composed of irradiated murine fibroblasts, that are essential for keratinocyte growth.^[8]^ The use of CEA introduces potential complications, such as xenogeneic contamination along with regulatory and ethical concerns.^[9,10]^ These limitations underscore the need for alternative scaffold systems that support cell adhesion and proliferation, in a stable and cost-effective manner, while avoiding the dependency on animal-derived components.

Peptide self-assembly is a promising strategy for skin wound healing and regeneration, utilizing short peptide sequences that form nanoscale structures resembling the extracellular matrix (ECM).^[11,12]^ Short peptides and amino acids can be chemically modified^[13]^, co-assembled^[14,15]^, combined with polysaccharides^[16]^ and incorporated with antibacterial and bioactive motifs.^[17–19]^ One prominent bioactive motif is the tripeptide arginine-glycine-aspartic acid (RGD). This sequence is a well-recognized integrin-binding motif, that plays a crucial role in promoting cell adhesion, signaling, and tissue regeneration.^[20,21]^ RGD facilitates the attachment of cells, such as fibroblasts and keratinocytes, to the ECM by binding to integrin receptors on the cell surface.^[22]^ This interaction anchors the cells to the scaffold and triggers intracellular signaling pathways that regulate cellular proliferation, migration, and differentiation.^[23]^ The interaction of fibroblast, keratinocyte, and endothelial cell integrins with the RGD motifs within fibronectin plays a crucial role in wound healing processes, including wound contraction, re-epithelialization, and angiogenesis, respectively.^[23]^

In addition, the fluorenylmethoxycarbonyl-diphenylalanine (Fmoc-FF) hydrogel self-assembled to form a stable, biocompatible scaffolds that provide structural support while allowing for controlled biodegradability.^[24–26]^ Under physiological conditions, Fmoc-FF efficiently self-assembles into a rigid fibrous hydrogel with a 3D ECM-like nanostructure.^[27–29]^ The Fmoc-FRGD peptide combines the self-assembling properties of the Fmoc-FF core with the bioactivity of the RGD motif.^[30]^ This unique composition results in a scaffold with enhanced biocompatibility, and bioactivity, creating an optimal environment for cell growth and tissue repair.

In recent years, tissue engineering has made significant advances, particularly in the design of electrospun polymeric scaffolds for advanced wound dressings, especially for severe burns.^[31,32]^ These scaffolds closely replicate the natural extracellular matrix (ECM), offering a large surface area and high porosity that create an optimal microenvironment for cellular attachment, proliferation, and differentiation.^[33]^ Despite their promise, many existing polymer nanofiber scaffolds used in wound healing suffer from limitations such as brittle or plastic mechanical behavior and low bioactivity, which hinder their effectiveness.^[34]^

Among synthetic polymers, poly(ε-caprolactone) (PCL) is widely regarded as an ideal material for electrospinning due to its excellent biocompatibility, solubility, and ability to blend with other materials. As an FDA-approved polyester, PCL exhibits long-term mechanical and structural stability, making it particularly suitable for applications requiring extended tissue growth and remodeling, such as chronic wound healing.^[35–37]^

To further enhance the functionality of electrospun scaffolds, the incorporation of bioactive agents has become a key strategy in tissue engineering.^[37,38]^ These agents include natural bioactive compounds, growth factors, enzymes, peptides, and other biomolecules, each tailored to address specific therapeutic needs. For instance, natural bioactive compounds such as plant extracts and oils have been investigated for their wound and burn healing potential but are often limited by challenges like chemical instability, poor solubility, oxidative degradation, and low bioavailability.^[39]^ Similarly, growth factors like bone morphogenetic proteins (BMPs) and vascular endothelial growth factor (VEGF) have been integrated into electrospun fibers to promote cell differentiation and tissue vascularization. However, their rapid degradation and loss of bioactivity necessitate protective encapsulation or controlled-release mechanisms to ensure sustained therapeutic effects.^[40]^ Enzymes like collagenase have also been incorporated into electrospun fibers to mimic natural tissue remodeling by modifying the ECM, but their integration requires careful control to prevent unintended degradation or loss of activity.^[41]^ Another promising approach involves the incorporation of molecular recognition sequences, such as RGD and its derivatives, which facilitate cell attachment and regulate cellular behaviors such as proliferation, migration, and differentiation. While these sequences significantly enhance scaffold bioactivity, their stability within the scaffold matrix and controlled release remain challenging, necessitating precise optimization of their loading and release profiles.^[42]^ Building on these advancements, our recent work has demonstrated the potential of incorporating bioactive peptides, specifically Fmoc-FRGD, into PCL electrospun scaffolds to improve bone regeneration.^[43]^ This peptide enhances cell adhesion by interacting with integrins and supports the development of functional tissues by promoting specific cellular responses. Unlike other bioactive systems, Fmoc-FRGD peptides offer a stable, bioactive structure that enables controlled cell-material interactions, addressing challenges such as instability or uncontrolled release seen in other peptide-based approaches. Our findings indicate that PCL scaffolds functionalized with Fmoc-FRGD exhibit superior biological performance, including enhanced cell attachment and improved integration with surrounding tissues in vivo, positioning them as a promising candidate for skin grafts and skin regeneration following burn injuries.

Here, we introduce a novel approach to wound healing through the development of a fully autologous skin equivalent system, integrating patient-derived cells within an advanced electrospun scaffold (Figure 1). The scaffold combines polycaprolactone (PCL) with Fmoc-FRGD peptide sequence, strategically designed to overcome critical limitations in current wound healing therapies. Through comprehensive *in-vitro* and *in-vivo* evaluations, our system demonstrates remarkable advantages: it maintains complete autologous properties, exhibits no shrinkage, provides superior mechanical stability, and significantly enhances cellular integration and wound healing. The scaffold’s unique composition promotes optimal tissue regeneration while maintaining practical considerations for clinical translation. These characteristics, combined with its scalability and user-friendly application, position this system as a promising therapeutic solution for advanced wound care and tissue regeneration applications.

**Figure 1.**
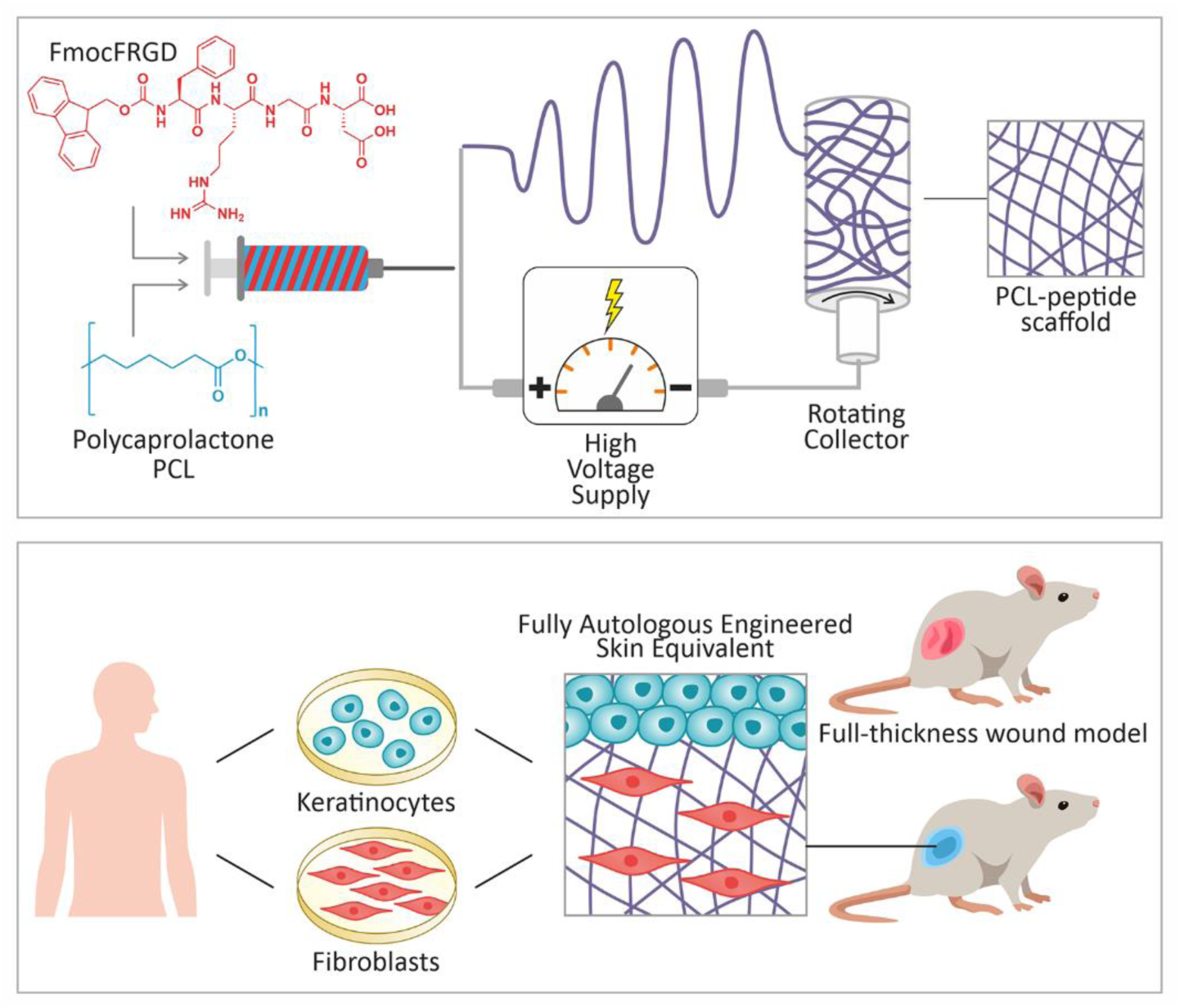
Schematic illustration of the development of a fully autologous skin equivalent system, integrating patient-derived cells within an advanced electrospun scaffold for the treatment of severe burn injuries. Design and fabrication of nanofiber scaffolds through electrospinning nonwoven nanofiber meshes using a single needle to produce uniform fibers (Upper panel). Isolation of human dermal fibroblasts and keratinocytes from patient biopsies to engineer the fully autologous skin equivalent scaffold, tested in a full thickness wound model (Lower Panel).

## Results and Discussion

### Physicochemical Structure and Hydrophilicity Characterizations of the PCL-peptide Nanofibers

The nanofiber scaffolds were designed and prepared by electrospinning nonwoven nanofiber meshes, as previously reported.^[43]^ Three distinct fiber compositions were examined: (a) pure PCL fibers, (b) PCL fibers with a 2.5% Fmoc-FRGD peptide-to-PCL ratio, and (c) PCL fibers with a 20% Fmoc-FRGD peptide-to-PCL ratio. This design maintained consistent polymer content while varying peptide concentrations across low and high levels to assess the influence of RGD functionalization on fiber properties and potential applications. In a typical procedure, the components were mixed, dissolved and dispensed through a single needle to obtain single component fibers.

Figure 1 presents the surface morphology, nanostructure, chemical structure and hydrophilicity of the PCL-Peptide nanofiber scaffolds. The morphology of the PCL-peptide nanofibers was characterized using high-resolution scanning electron microscopy (HR-SEM) (Figure 1a). The micrographs revealed that the nanofiber matrix has a three-dimensional porous structure that closely mimics the architecture of the ECM.^[44]^ This biomimetic porosity is essential for supporting cell infiltration, nutrient diffusion, and waste removal.^[45]^ The nanofiber diameters of pure PCL and PCL-peptide were determined as 299 ± 64 nm (PCL); 332 ± 48 nm (PCL-2.5%Fmoc-FRGD); 369 ± 51 nm (PCL-20%Fmoc-FRGD) (Figure 1b). Thus, the incorporation of the Fmoc-FRGD peptide slightly increased the nanofiber diameter of PCL.

Fourier transform infrared (FTIR) spectroscopy analysis provided insights into the chemical structure of PCL/Fmoc-FRGD hybrid nanofibers (Figure S1). Characteristic C–H stretching vibrations from methylene groups in the PCL backbone were observed at 2943 and 2865 cm^-1^, carbonyl stretching (C=O) appeared at 1721 cm^-1^, and −C−O−C− absorption peaks were detected at 1293 and 1240 cm^-1^ across all samples. Notably, in the PCL/Fmoc-FRGD samples, additional peaks were detected at 1660 cm^-1^ and 1540 cm^-1^ corresponding to the amide I and amide II bands, respectively, associated with the peptide backbone and indicating β-conformations. Combined with minor peaks at 1674 and 1689 cm⁻¹, these peaks further support the presence of β-sheet secondary structures within the peptide component.^[30,46]^ The peak at 1540 cm^-1^ also aligns with the aromatic ring vibrations of the Fmoc group, confirming the successful incorporation of Fmoc-FRGD in the nanofiber structure.^[47]^

Controlled hydration is crucial for biomaterials utilizes for wound healing.^[48]^ Figure 1c-g illustrates the hydrophilicity and water absorption characteristics of the PCL−Peptide nanofibrous scaffolds. The contact angle of the pure PCL nanofiber scaffold was measured to be 98.46° ± 0.7° and remained stable over 4-minutes (Figure 1c, e, g). In contrast, the contact angles of the PCL-Peptide scaffolds were initially higher, measuring 126.03° ± 0.54° for PCL-2.5% Fmoc-FRGD and 122.89° ± 2.68° for PCL-20% Fmoc-FRGD. However, after 4 minutes, these angles decreased significantly to 50.78° ± 4.48° and 41.24° ± 0.14°, respectively (Figure 1e, g). This reduction is likely attributed to the hydrophilic nature of the Fmoc-FRGD peptides embedded in the scaffold, which emanates from the arginine (R) and aspartic acid (D) hydrophilic amino acids that have charged side chains, making this part of the peptide capable of strong interactions with water. To further enhance hydrophilicity, the nanofibrous scaffolds underwent oxygen plasma treatment, which introduces polar functional groups on the surface and is a widely recognized method for improving the hydrophilic properties of materials.^[48,49]^ The plasma treatment was applied for 45 seconds at low intensity, significantly increasing hydrophilicity (Figure 1d, f, g). Following plasma treatment, the contact angle of the pure PCL nanofiber scaffold decreased to 40.1° ± 0.62° and further reduced to 22.06° ± 4.3° after 4 minutes. The contact angles for the PCL-Peptide scaffolds were measured to be 21.68° ± 0.28° for PCL-2.5% Fmoc-FRGD and 23.05° ± 0.05° for PCL-20% Fmoc-FRGD, with both showing a dramatic decrease to 0° within the same timeframe (Figure 1f, g). This decrease can also be attributed to capillary effects that enhance the penetration of the liquid into the porous matrix. Thus, we can conclude that the combination of peptide incorporation and plasma treatment resulted in a highly hydrophilic matrix, potentially enhancing its ability to facilitate the transport of nutrients and manage extravasation during the wound healing process.

**Figure 1.**
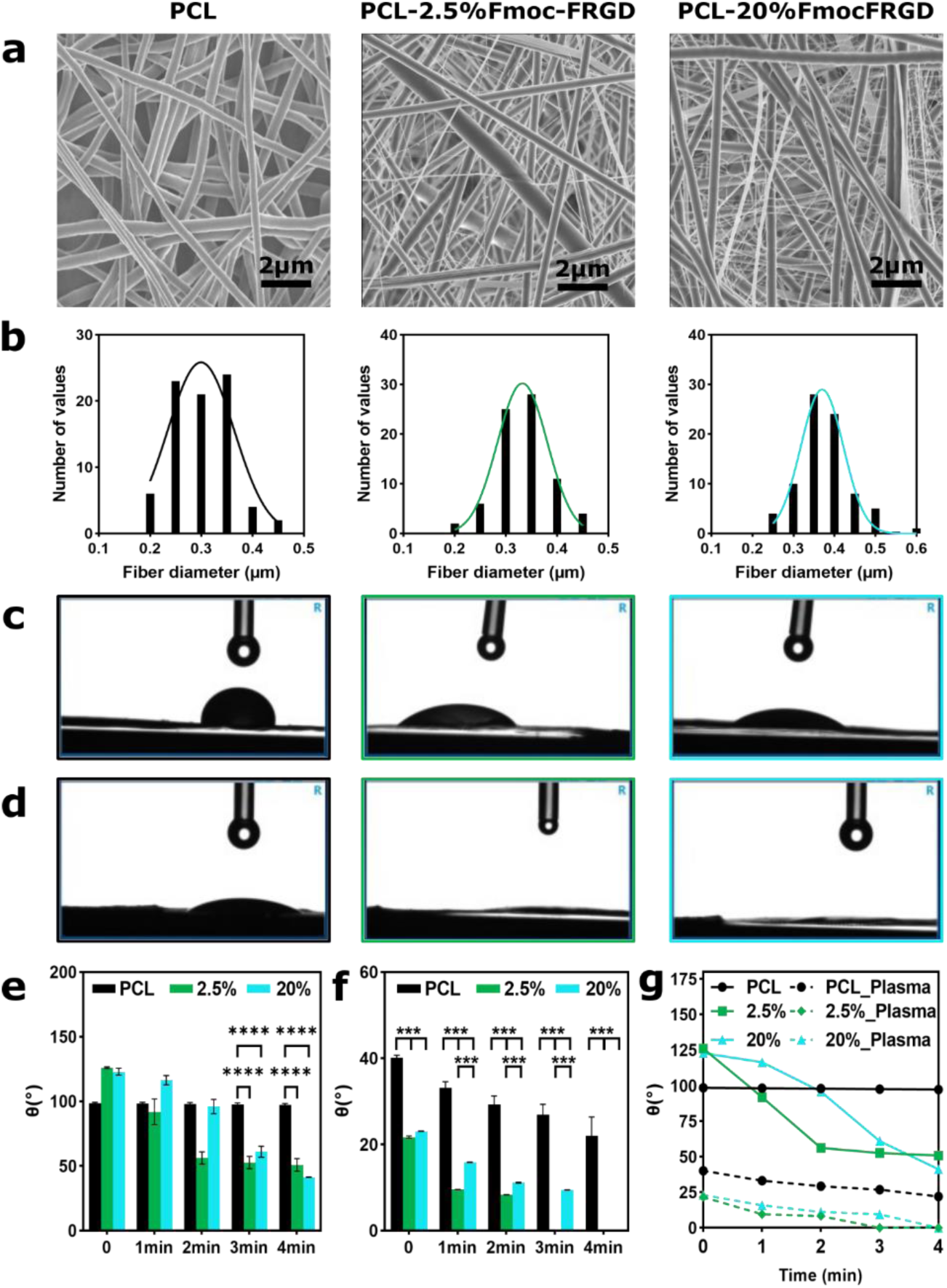
Characterizations and hydrophilicity analysis of the PCL−Peptide hybrid nanofibrous scaffolds. (a) HR-SEM images of the PCL and PCL−Peptide nanofibrous matrix. (b) Diameter distribution of the nanofibers. (c-d) Water drops change on the different matrices (c) before and (d) after plasma treatment. (e-g) Water contact angle test for 4 min (e) before and (f) after plasma treatment. Statistical significances are indicated as ***p < 0.001 and ****p < 0.0001.

### Biomimetic Mechanical Properties of the PCL-Peptide Nanofibers

Analyzing the mechanical properties of electrospun scaffolds is vital for their application in tissue engineering and personalized skin grafts. These properties dictate the scaffold’s structural integrity, enabling it to withstand physiological forces while mimicking the behavior of native skin. Moreover, optimizing the mechanical properties (e.g. flexibility and stiffness) allows for customization based on individual patient needs, ultimately enhancing healing outcomes and ensuring scaffold durability.

The mechanical properties of electrospun PCL and PCL-peptide nanofibers were evaluated using uniaxial tensile testing. Figure 2a presents the stress-strain curves for scaffolds composed of pure PCL, PCL-2.5%Fmoc-FRGD, and PCL-20%Fmoc-FRGD, all with comparable fiber densities and dimensions.

**Figure 2.**
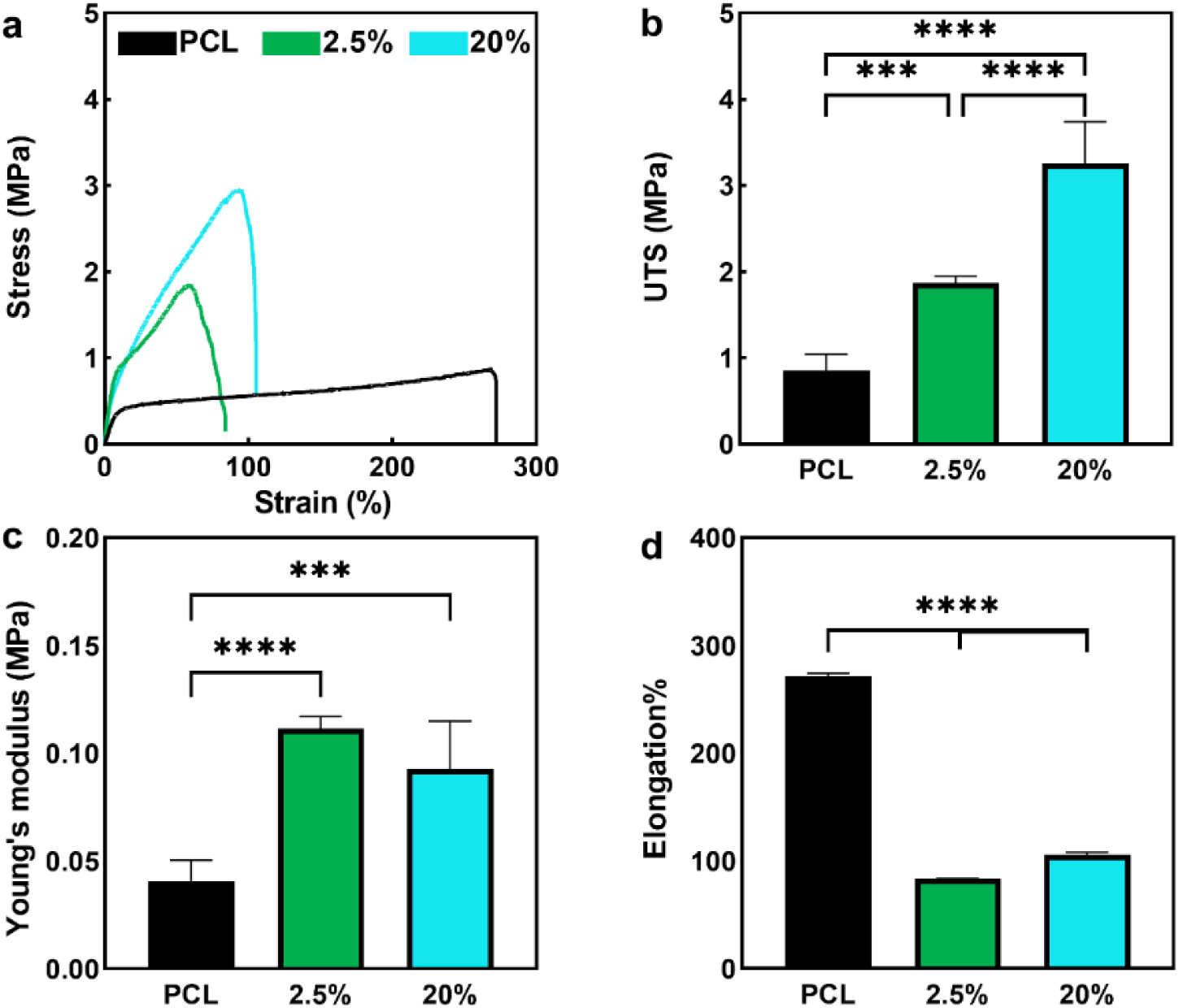
Biomimetic Mechanical Properties of the PCL-peptide Nanofibers. (a) Stress−strain curves of the scaffolds. (b) Ultimate tensile strength. (c) Young’s modulus. (d) Elongation at the break. Statistical significances are indicated as ***p < 0.001 and ****p < 0.0001.

The corresponding ultimate tensile strength (UTS) values are presented in Figure 2b. Notably, the PCL-20%Fmoc-FRGD group demonstrated the highest UTS, significantly surpassing both pure PCL and PCL-2.5%Fmoc-FRGD. This trend highlights a clear enhancement in UTS with increasing peptide concentration, suggesting that peptide incorporation strengthens the fiber’s structural integrity through enhanced intermolecular interactions between the peptide and the polymer matrix.^[50,51]^

Young’s modulus of the electrospun fibrous scaffolds was evaluated to assess the stiffness of the scaffolds (Figure 2c). Pure PCL exhibited the lowest Young’s modulus of 40.6 ± 9.7 kPa, reflecting its relatively high flexibility and lower resistance to deformation. The incorporation of 2.5%Fmoc-FRGD peptide led to a substantial increase in stiffness, with the highest modulus of 111.6 ± 5.5 kPa, suggesting enhanced fiber rigidity and stronger intermolecular bonding.

Interestingly, the 20%Fmoc-FRGD fibers, despite their superior UTS, exhibited a slightly lower mean Young’s modulus of 92.75 ± 22.0 kPa, indicating that at higher peptide concentrations, the material becomes more deformable, potentially due to changes in fiber microstructure. When compared to native skin tissue, which typically has a tensile strength of 1-30 MPa,^[52]^ the PCL-20%FmocFRGD nanofibers exhibit a comparable mechanical profile, positioning this scaffold as a promising candidate for skin regeneration.

The tensile displacement provides critical insight into the elongation capacity of the electrospun fibers before failure (Figure 2d). The pure PCL fibers exhibited the highest mean tensile displacement of 271% ± 2.9%, reflecting their significant flexibility and ability to stretch underload. In contrast, the incorporation of Fmoc-FRGD peptide resulted in a marked reduction in tensile displacement, with the PCL-2.5%Fmoc-FRGD nanofibers displaying a mean value of 83% ± 0.6% tensile displacement and the PCL-20%Fmoc-FRGD nanofibers showing 106% ± 1.9% tensile displacement. This substantial decrease in elongation upon peptide incorporation suggests that while the modified fibers offer enhanced stiffness and tensile strength, they become less capable of undergoing large deformations. The reduced displacement in PCL-2.5%Fmoc-FRGD aligns with its increased stiffness, indicating a trade-off between flexibility and structural reinforcement.

The stiffness displayed by the scaffold aligns well with the mechanical requirements of soft tissues, particularly skin, basal membrane, and subcutaneous connective tissues.^[53]^ Native skin typically has Young’s modulus ranging from 0.42 to 0.85 MPa depending on its location and depth^[54]^, with the dermis displaying higher moduli due to collagen content. The PCL-peptide scaffolds are softer but still fall within the lower range of required mechanical properties. This suggests they are suitable for skin graft applications, where both flexibility and strength are essential to mimic the dynamic environment of human skin. Furthermore, the stiffness of these scaffolds exceeds that of many conventional wound dressings, which prioritize flexibility over strength.

Interestingly, the peptide-modified scaffolds exhibit mechanical properties that mimic the structural role of a basement membrane, which has an approximate modulus of 95-100 kPa^[55]^, providing a stable framework for tissue integration. This balance between stiffness and flexibility is critical for scaffolds aimed at serving as personalized skin grafts, where both mechanical strength and the ability to conform to the body’s dynamic environment are essential. Among the three groups examined, the PCL-2.5%Fmoc-FRGD scaffold emerges as the most suitable candidate for skin grafting applications due to its balanced mechanical properties. This group exhibited the highest Young’s modulus (111.6 ± 5.5 kPa), indicating enhanced stiffness compared to the pure PCL and the PCL-20%Fmoc-FRGD, while maintaining sufficient flexibility to support dynamic mechanical environments (83% ± 0.6% tensile displacement). The elevated ultimate tensile strength (1.86 MPa) further highlights its capacity to withstand mechanical stress, surpassing pure PCL fibers, and closely aligned with the lower range of tensile properties observed in native skin.

### Contraction Behavior of the PCL-peptide Nanofibers Compared to CEA

A major challenge with CEAs is their pronounced shrinkage upon detachment from the culture plate, which complicates production processes and reduces overall yield. As shown in Figure 3a, the CEA retains its full size after 14 days of growth in culture dish. However, following detachment using Dispase II, a mild enzyme that cleaves proteins at the dermal-epidermal junction to gently release the cell sheet, the CEA rapidly contracts. Figure 3b illustrates this process, where the CEA shrinks to 21.62% ± 0.72% of its original size, even before transplantation. This well-defined contraction, occurring immediately after detachment, reduces the effective surface area available for grafting, presenting a substantial limitation. Addressing this contraction behavior is critical for improving the stability and usability of CEA sheets in clinical applications. The contraction of CEA sheets is primarily caused by the lack of a dermal component and the contractile properties of the keratinocytes, which cause the sheet to pull in upon detachment. Additionally, the CEA sheets cannot be directly handled for transplantation in their detached state. To lift and transfer the CEA sheets to the wound site, there is a need for additional tools, such as gauze pads, to support and stabilize the sheet during the procedure. This complicates the graft preparation process, making it more challenging to handle. In contrast to CEA, the electrospun fibers developed in this study exhibited no contraction throughout the entire process, maintaining their original dimensions both after fabrication and during cell culture and application (Figure 3c-f). As shown in Figure 3c, the electrospun nanofiber scaffold was cut into rectangular sections and placed in a culture dish, demonstrating its ease of handling and adaptability to custom shapes as required for specific applications. These scaffolds are easy to handle, as they can be securely grasped with tweezers and moved around without difficulty (Figure 3d). After cell culture, the electrospun nanofiber scaffold could be easily removed from the culture medium using tweezers (Figure 3e). Notably, as shown in Figure 3f, the electrospun nanofiber scaffold retained its original rectangular shape post-culture, with no signs of contraction or deformation. This structural stability, coupled with their robust mechanical properties, makes the electrospun fibers a superior alternative for ensuring consistent wound coverage and supporting tissue regeneration, without the challenges associated with CEA contraction.

**Figure 3.**
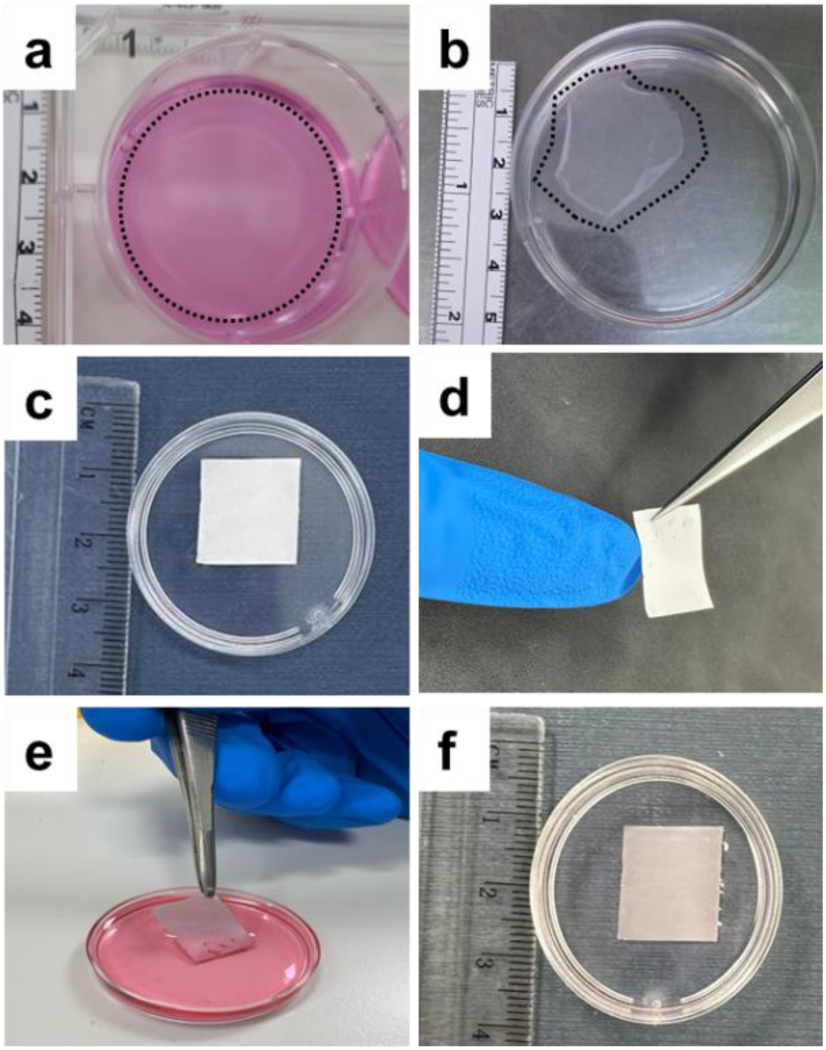
Contraction behavior of the CEA and the electrospun nanofibrous. (a-b) CEA after 14 days of culture (a) before and (b) upon detachment from the culture dish. The black dotted line circles the different samples sizes. (c) The electrospun nanofibrous scaffold is cut into a rectangular shape. (d) The scaffold could be easily handled and held with tweezers. (e) The scaffold was soaked in cell media and easily pulled out with tweezers. (f) The scaffold after cell culture.

### Human Dermal Fibroblast and Keratinocyte Cells Seeding on the PCL-peptide Scaffolds

Human dermal fibroblasts and keratinocytes were isolated from patient biopsies, expanded *ex-vivo*, and directly seeded onto the scaffolds—PCL, PCL-2.5%Fmoc-FRGD, and PCL-20%Fmoc-FRGD—without the need for a feeder layer. The biocompatibility of the PCL/peptide scaffolds was assessed *in-vitro* at multiple time points (days 1, 4, 8, and 11) using the XTT assay (2,3-bis-(2-methoxy-4-nitro-5-sulfophenyl)-2H-tetrazolium-5-carboxanilide), as shown in Figure 4a. The results revealed notable differences in cell viability among the three experimental groups. By day 4, the PCL-2.5%Fmoc-FRGD scaffold exhibited significantly higher cell viability compared to both the PCL and the PCL-20%Fmoc-FRGD. This trend continued through days 8 and 11, with a slight increase in cell viability on the PCL scaffold, while both Fmoc-FRGD groups maintained substantially higher viability, highlighting the enhanced bioactivity of the peptide-modified scaffolds.

**Figure 4.**
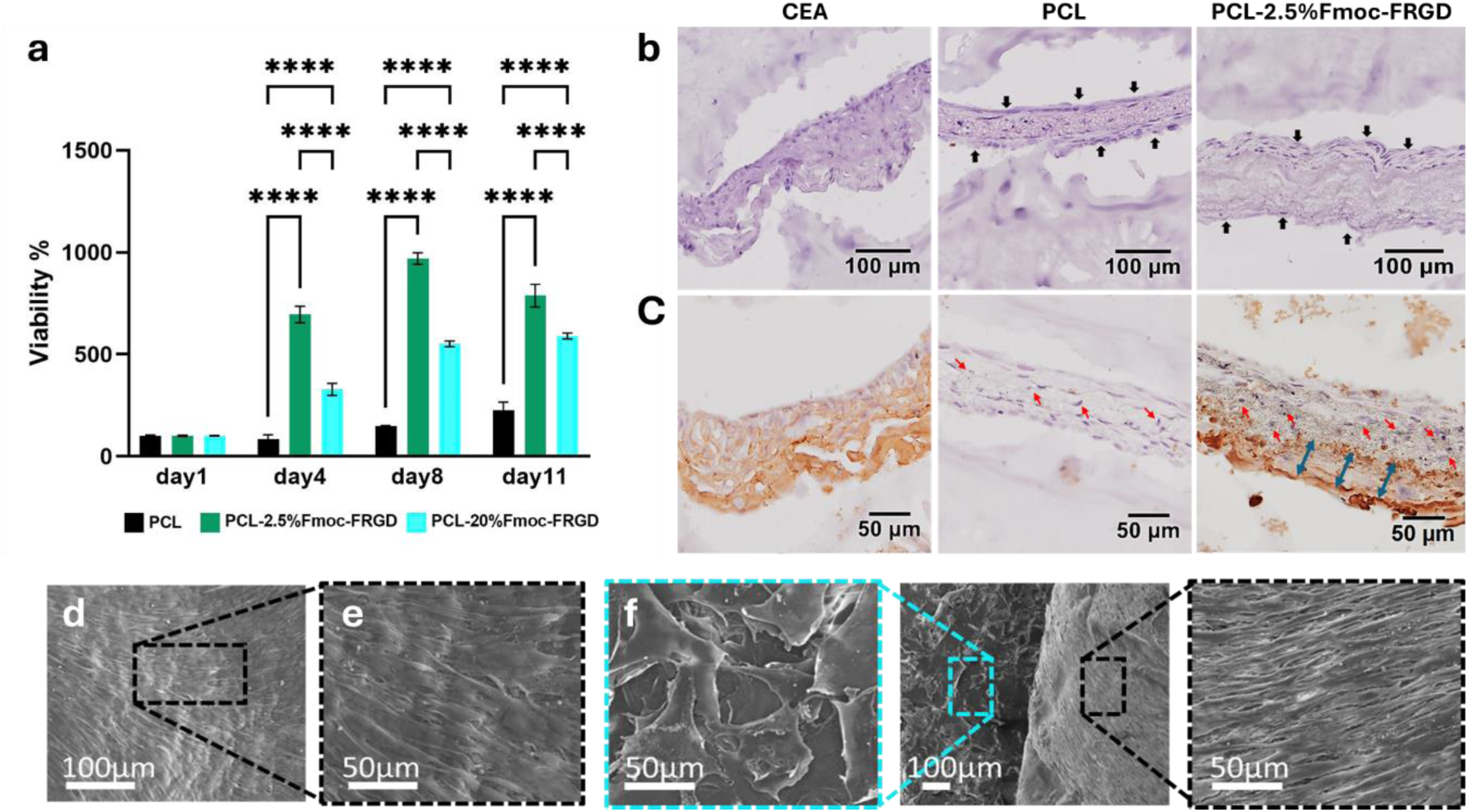
Human Dermal Fibroblast and Keratinocyte Cells Seeding on the PCL-peptide Scaffolds. (a) Cell viability analysis of mixed fibroblast and keratinocyte cells derived from a human skin biopsy at different time points. (b-c) Histological cross-sections of the CEA, PCL and the PCL-2.5%Fmoc-FRGD following cell culture, evaluated using H&E staining. Black and red arrows indicate cell attachment and infiltration into the scaffolds respectively. (b) and cytokeratin (CK10) immunostaining. Blue arrows indicate the presence of keratinocytes. (c). (d-f) HR-SEM images of the PCL−peptide scaffold after cell culture, (d-e) The PCL scaffold showing growth and proliferation of fibroblasts. (f) The PCL-2.5%Fmoc-FRGD scaffold showing the growth and proliferation of keratinocyte on one side (left) and fibroblast on the other side (right). Statistical significances are indicated as ****p < 0.0001.

Notably, the PCL-2.5%Fmoc-FRGD group exhibited the highest levels of cell proliferation, maintaining superior viability throughout the culture period. This analysis demonstrated that the electrospun fibers provided an optimal surface for cell growth, enabling proliferation directly on the scaffold without the need for a murine-derived feeder layer. This offers a significant advantage for skin grafts by reducing preparation time, as it bypasses the feeder layer step typically required for culturing CEA sheets, potentially accelerating clinical application and improving regulatory efficiency.

These results indicate that the PCL/Fmoc-FRGD scaffolds significantly promote cell proliferation, particularly at lower peptide concentrations (2.5%). The highest levels of cell viability observed for the PCL-2.5%Fmoc-FRGD fibers suggest that this moderate concentration of the bioactive peptide provides an optimal environment for cell attachment and proliferation, enhancing the scaffold’s biocompatibility. In contrast, while the PCL-20%Fmoc-FRGD group also shows improved cell viability compared to the pure PCL fibers, the enhancement is less pronounced than that observed for the 2.5% group. This suggests that higher concentrations of the peptide may lead to a saturation effect or changes in the material properties of the scaffold, which could limit its effectiveness.^[56,57]^ Based on these results, the PCL-2.5%Fmoc-FRGD scaffold was chosen for further study, as it offered the best balance of biocompatibility and functionality.

Mixed human-derived skin cells were seeded on the scaffolds and cultured until the end of the healing period, aligning with the timeframe of the *in-vivo* experiments. To gain deeper insights into cellular adhesion, infiltration, and organization within the CAE and the electrospun scaffolds, we performed histochemical analyses. Hematoxylin and eosin (H&E) staining was employed to evaluate tissue morphology (Figure 4b), and anti-cytokeratin (CK10) immunostaining was used to identify keratinocytes specifically (Figure 4c).

The CEA (left panels) exhibited a fragile structure, reflecting its inherent limitations. Comprised solely of epidermal cells (Figure 4c, left), the CEA lacked the essential mechanical stability and three-dimensional architecture required to support sustained cellular integration and coordinated dermal-epidermal organization.

The PCL scaffold (middle panels) showed evidence of cellular attachment and limited infiltration, with cells primarily aligned along the scaffold’s periphery, as indicated by the black arrows in Figure 4b. Despite some fibroblast infiltration (red arrows), the overall cell distribution lacked depth and organization, and no evidence of keratinocyte was detected in CK10 staining. This suggests that the pure PCL scaffold lacks the necessary surface chemistry or bioactive cues to support the formation of stratified, well-organized skin-like tissue.

Notably, the peptide-functionalized PCL-2.5%Fmoc-FRGD scaffold (right panels) demonstrated significantly enhanced cellular adhesion, infiltration, and multi-layered organization, closely mimicking native skin tissue architecture. H&E staining revealed a dense cell population on both the surface and within the scaffold (black and red arrows, respectively), while CK10 staining (blue arrows) confirmed the presence of keratinocytes, highlighting its capacity to support epidermal differentiation. We propose that the mechanism for spontaneous dermal-epidermal organization in the PCL-2.5%Fmoc-FRGD scaffold is attributed to the synergistic interplay of its biochemical, physical, mechanical, and microenvironmental properties.^[58,59]^ The incorporation of Fmoc-FRGD introduces RGD motifs that bind integrins on fibroblasts and keratinocytes, enabling selective adhesion and spatial organization. Fibroblasts infiltrate the scaffold’s porous nanofibrous structure, and can produce ECM components like collagen and fibronectin, while the scaffold’s hydrophilic surface supports keratinocyte adhesion and stratification on the surface. The scaffold’s ECM-like architecture further guides cells spatially, mimicking natural tissue organization.^[33]^ Additionally, it is known that paracrine signaling between fibroblasts and keratinocytes enhances their alignment into distinct dermal and epidermal layers, recreating the cues necessary for tissue regeneration without external stratification techniques.^[60]^

Previous studies have shown that a highly stiff substrate (Young’s modulus ≈ 200 kPa) promotes epidermal cell proliferation, migration, and re-epithelialization.^[61]^ In this context, the mechanical properties and stiffness of the PCL-2.5%Fmoc-FRGD scaffold (Young’s modulus ≈ 111.6 ± 5.5 kPa) provide an optimal balance of substrate tension and mechanical cues, supporting fibroblast and keratinocyte adhesion, proliferation, and migration. These properties facilitate the formation of a functional, organized skin equivalent in vitro, demonstrating the scaffold’s potential in advanced tissue engineering applications.^[61,62]^

HR-SEM images confirmed the distribution and attachment of cells on the two different scaffolds. Figures 4d and 4e illustrate how fibroblasts adhered to the surface of the PCL scaffold, clearly demonstrating their attachment and spreading across the fibers. Figure 4f provides a detailed view of both sides of the PCL-2.5%Fmoc-FRGD scaffold simultaneously, showing distinct cell behavior on each side. On one side, aligned fibroblasts were observed, forming a well-organized dermal layer (Figure 4f, right), while on the opposite side, keratinocytes were detected, forming an epidermal layer (Figure 4f, left). This dual-layer cell growth highlights the scaffold’s ability to promote organized, full-thickness skin equivalent without the need for external patterning or stratification.

The formation of these distinct cell layers within the PCL-2.5%Fmoc-FRGD scaffold is a significant advancement since full-thickness skin consists of both dermal (fibroblast-rich) and epidermal (keratinocyte-rich) layers, which are essential for its function and resilience. The PCL-2.5%Fmoc-FRGD scaffold not only supports the growth of both cell types but also facilitates their spatial organization, creating a more robust and functional skin equivalent. This organized, full-thickness structure is critical for skin grafts, as it provides both the strength, and the biological function required for integration and repair in wound sites.

### *In-vivo* Wound Healing Analysis

The wound healing efficiency of the engineered skin equivalents was evaluated in an in vivo model using nude mice. These mice, genetically engineered to lack a functional thymus, have a suppressed immune system with significantly reduced T-cell levels, enabling the implantation of human cells without immune rejection. Full-thickness wounds (1 cm²) were created on the backs of the mice, then treated with different scaffold types: untreated control, CEA, engineered skin equivalents based on the PCL or PCL-2.5%Fmoc-FRGD scaffolds populated with human skin cells (Figure 5). Wound closure progress was recorded through photographs and quantified measurements. Over a 19-day period, the PCL-2.5%Fmoc-FRGD scaffold demonstrated the most rapid wound closure, significantly outperforming both untreated control and CEA. The wound area measurements (Figure 5b-c) showed that by day 4, the PCL-2.5%Fmoc-FRGD group had reduced wound size by 45.9% ± 15.7%, while untreated, CEA-treated and PCL, groups displayed slower reduction rates of 16.2% ± 2.2%, 33.0% ± 9.1%, and 36.9% ± 1.3%, respectively. By day 5, 50% of the wound was closed in the PCL-2.5%Fmoc-FRGD group. In comparison, it took 7 days to reach this level of closure in the PCL group, 8 days in the CEA group, and up to 12 days in the untreated control group. This trend proceeded on day 8, where all treatment groups, CEA, PCL, and PCL-2.5%Fmoc-FRGD exhibited significantly better wound closure compared to the untreated group (\*\*\*\**, p < 0.0001).* Notably, the PCL-2.5%Fmoc-FRGD group achieved better wound closure than the commonly used CEA. By day 19, wounds covered with the PCL-2.5%Fmoc-FRGD skin equivalent achieved nearly complete closure, with wound size reduced to just 0.05% of the original area, compared to 2.5% in untreated wounds and 0.33% in CEA-treated wounds. The PCL-2.5%Fmoc-FRGD skin equivalent demonstrated an accelerated wound healing rate, particularly in the early phases of wound closure (days 4–11), compared to the untreated and CEA groups. This early enhancement in wound healing may provide a crucial advantage for rapid tissue regeneration, reducing the time required for wound closure. This superior healing outcome may be attributed to the bioactive Fmoc-FRGD peptide, which enhances cell attachment and proliferation, creating a favorable environment for tissue regeneration.

**Figure 5.**
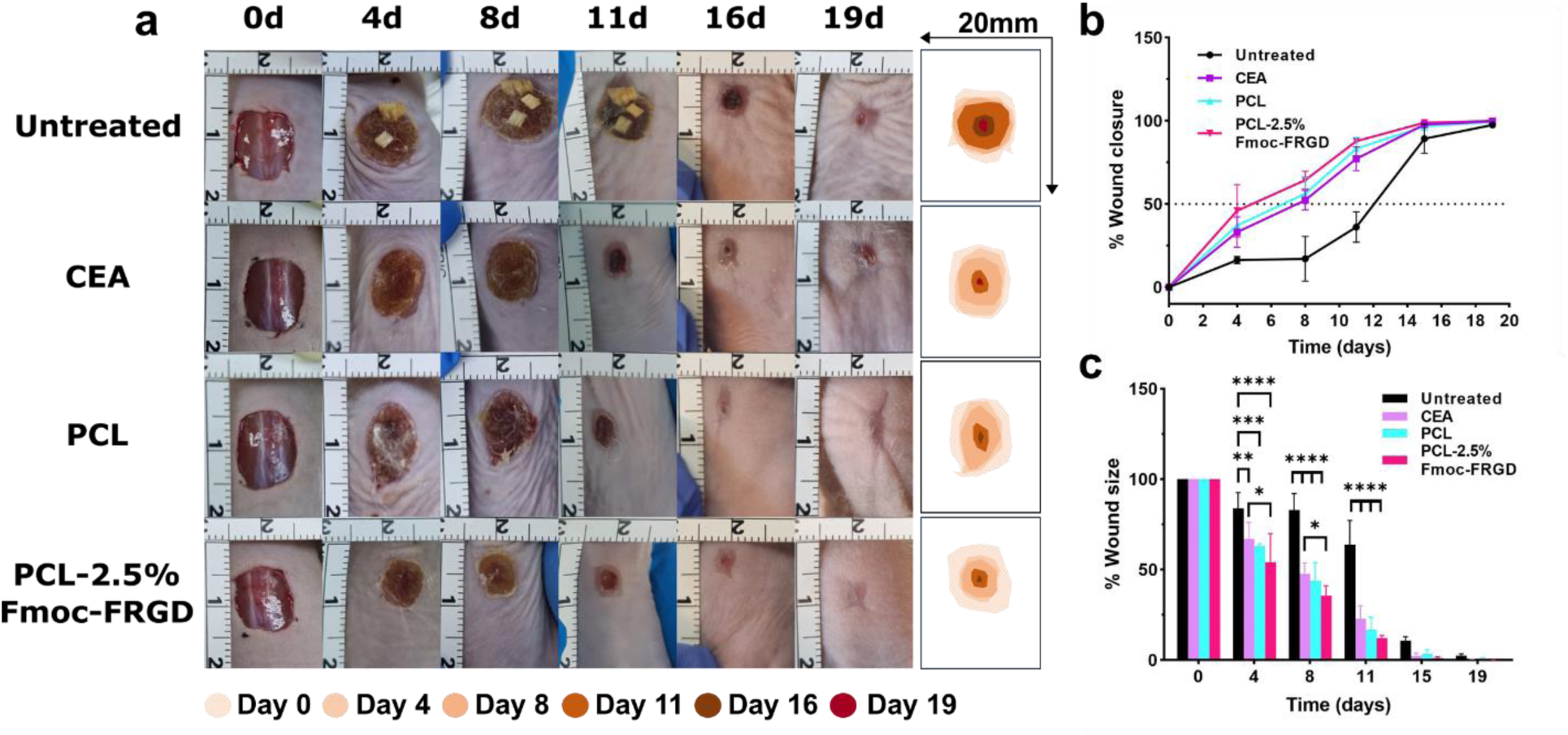
*In vivo* evaluation of the wound healing process. (a) Photograph of wounds untreated (control) and treated with CEA, PCL, PCL-Fmoc-FRGD2.5% for 0, 4, 8, 11, 16 and 19 d (scale bar = 20 mm). (b) Wound closure rate at different time points. The dotted black line marks 50% wound closure. (c) Quantification of wound size at different time points. Statistical significances are indicated as *p < 0.05, **p < 0.01, ***p < 0.001 and ****p < 0.0001.

Histological analysis using H&E staining revealed significant differences in tissue regeneration and organization across the treatment groups (Figure 6a). In the untreated control group, minimal epidermal regeneration was observed, with a relatively thin epithelial layer and sparse dermal organization. The CEA-treated wounds showed improved tissue thickness compared to the untreated control, though the dermal-epidermal junction remained relatively uniform without significant rete ridge formation. The PCL treatment resulted in moderate epithelialization with some enhancement in dermal organization, suggesting a supportive role in the healing process.

**Figure 6.**
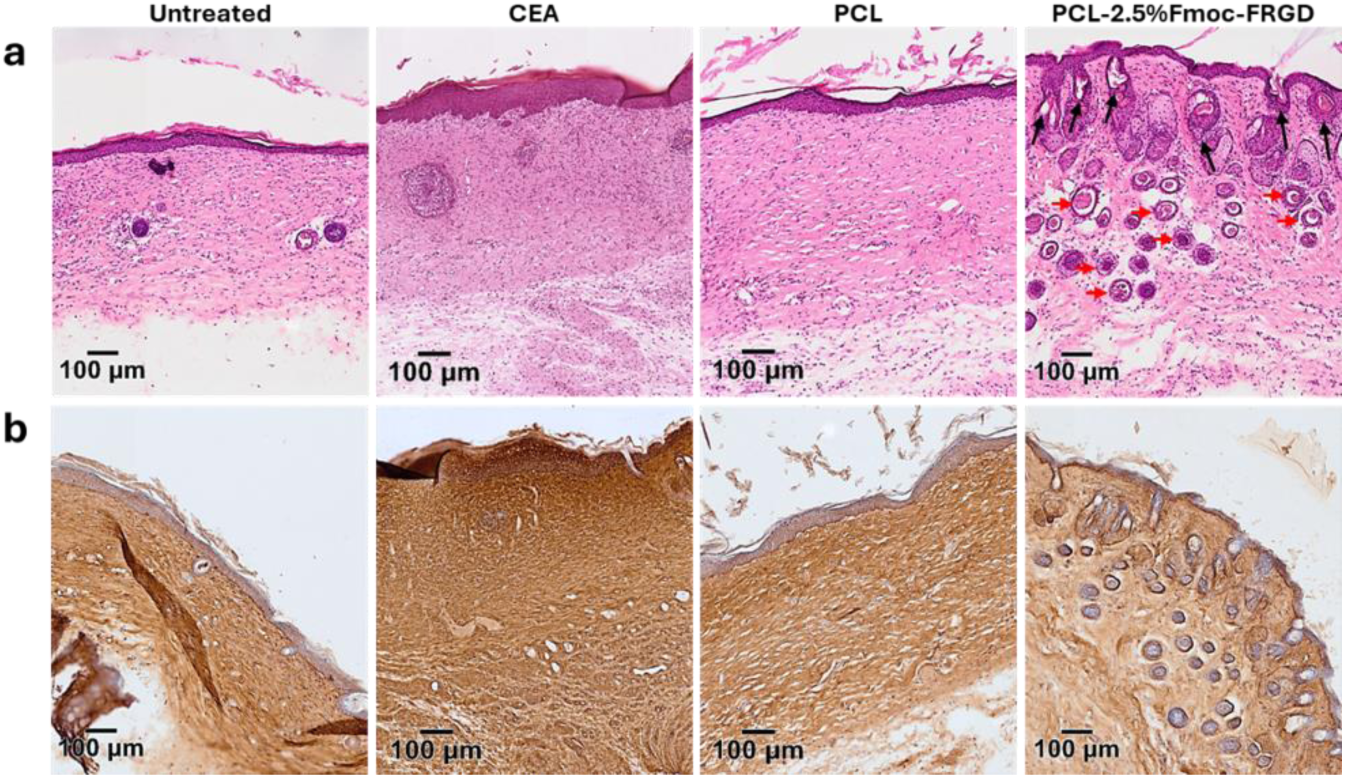
Histological evaluation of wound healing and skin regeneration *in-vivo*. (a) H&E staining of untreated wound tissues and treated with CEA, PCL or PCL-2.5%Fmoc-FRGD. Black and red arrows indicate hair follicles and secretory glands, respectively. (b) Immunohistochemistry CK10 staining of the wound tissues.

Notably, wounds treated with the PCL-2.5%Fmoc-FRGD exhibited exceptional healing characteristics, demonstrating advanced tissue regeneration at multiple levels. The treatment resulted in superior epidermal architecture characterized by pronounced epithelial stratification, distinct rete ridge formation, and a well-developed dermal-epidermal junction.^[63]^ Additionally, we observed the regeneration of skin appendages, with histological examination revealing the early-stage formation of hair follicles (black arrows) and secretory glands (red arrows) in the H&E staining, closely resembling those found in healthy mouse skin (Figure S2).^[64,65]^

These nascent appendages exhibited appropriate morphological organization and spatial distribution within the dermis, suggesting active tissue regeneration and maturation processes. The initiation of such complex structural formations is particularly noteworthy, as functional hair follicles and secretory glands are critical for maintaining skin homeostasis and physiological functions. The enhanced regenerative capacity of the scaffold likely results from the synergistic interplay between its mechanical properties and bioactivity, which together promotes the formation of the skin equivalent scaffold, as demonstrated in vitro. This combination forms an optimal microenvironment that supports not only initial cellular attachment and proliferation but also tissue maturation and functional development *in-vivo*.

Further studies using animal models that better replicate human skin regeneration—such as porcine burn models or other large-animal systems—will be essential to validate the potential of the PCL-2.5%Fmoc-FRGD skin equivalent scaffold in promoting complete and functional skin restoration in humans. Nevertheless, this comprehensive regeneration of both tissue architecture and appendages highlights the superior therapeutic potential of the PCL-2.5%Fmoc-FRGD scaffold in driving complete and functional skin restoration.

CK10 immunostaining provided further insights into keratinocyte differentiation and epidermal maturation (Figure 6b). The PCL-2.5%Fmoc-FRGD group demonstrated intense and uniform CK10 expression in the suprabasal layers, indicating proper terminal differentiation of keratinocytes and establishment of a mature epidermis. This enhanced expression pattern was particularly evident in the numerous rete ridges, suggesting improved mechanical stability and barrier function. In contrast, the untreated control showed minimal CK10 expression, while the CEA and the PCL groups displayed moderate CK10 staining patterns, which were less organized than the PCL-2.5%Fmoc-FRGD. These findings collectively suggest that PCL-2.5%Fmoc-FRGD treatment accelerates wound closure and promotes a more physiologically relevant tissue regeneration, characterized by proper stratification, differentiation, and structural organization of the regenerated skin tissue.

## Conclusions

The PCL-2.5%Fmoc-FRGD engineered skin equivalent demonstrates substantial advantages over conventional skin graft options, such as CEA. While CEAs have been established as a standard treatment for wound healing, they are limited by significant drawbacks including fragility, poor mechanical strength, and susceptibility to contracture. Our engineered PCL-2.5%Fmoc-FRGD skin equivalent effectively addresses these limitations while demonstrating superior performance across multiple critical parameters, positioning it as an optimal solution for severe burn treatment.

From a mechanical perspective, the PCL-2.5%Fmoc-FRGD skin equivalent exhibits remarkable stability and handling characteristics compared to both the CEA and the pure PCL scaffolds, enabling robust clinical application without compromising structural integrity. *In-vitro* studies reveal the scaffold’s capacity to support full-thickness skin development through spontaneous organization of fibroblasts and keratinocytes in physiologically relevant layers, closely mimicking natural skin architecture. This self-organizing capability eliminates the need for complex stratification procedures, significantly enhancing the scaffold’s clinical utility and effectiveness.

A key advancement of the PCL-2.5%Fmoc-FRGD system is the elimination of murine feeder layers, traditionally required in skin equivalent production. This improvement reduces potential contamination risks, simplifies regulatory compliance, and streamlines the manufacturing process.

*In-vivo* evaluation in a murine wound model revealed that the PCL-2.5%Fmoc-FRGD skin equivalent exhibited significantly enhanced wound healing compared to all other treatment groups. By day 5, wounds treated with the PCL-2.5%Fmoc-FRGD achieved 50% closure, markedly outperforming the CEA group (50% closure by day 7), the PCL group (day 8), and the untreated control (day 12). This rapid wound closure reduces the duration of wound exposure, thereby minimizing the risks of infection and systemic complications. Furthermore, the PCL-2.5%Fmoc-FRGD skin equivalent not only accelerates healing but also facilitates the regeneration of skin tissue with superior physiological relevance. The regenerated tissue demonstrated proper stratification, cellular differentiation, and structural organization, emphasizing the scaffold’s potential to support functional and durable skin restoration in clinical applications.

The PCL-2.5%Fmoc-FRGD skin equivalent represents a significant advancement in burn wound treatment, combining mechanical stability, ease of handling, and complete autologous properties with enhanced wound healing capabilities. Its superior performance in supporting organized tissue development, coupled with accelerated wound closure *in-vivo*, establishes this system as a promising therapeutic solution for severe burn injuries. These advantages, together with its practical clinical applicability, position the PCL-2.5%Fmoc-FRGD skin equivalent as an innovative approach to improving patient outcomes while minimizing the long-term complications associated with severe burns.

## Experimental Section/Methods

### Preparation of Electrospun fibers

For the electrospinning of PCL fibers (average Mn 80,000, Sigma-Aldrich (Rehovot, Israel)), 0.150 g of PCL was dissolved in 1.0 mL of hexafluoroisopropanol (HFIP) (Sigma-Aldrich (Rehovot, Israel)) (15% (w/v)), to obtain a clear homogeneous solution. For the electrospinning of PCL Fmoc-FRGD fibers, Fmoc-FRGD (GL Biochem (Shanghai, China)) was first dissolved in HFIP, at a concentration of 50 mg/ml. Next, the stock solution was added to the solution of the PCL in HFIP, described above. The solutions were combined to achieve a final composition of 15% (w/v) of PCL and 0, 0.375% and 3% (w/v) of Fmoc-FRGD, corresponding to a peptide-to-PCL ratio of 0:1, 1:40 (2.5%) and 1:5 (20%), respectively. Fresh stock solutions were prepared for each experiment, and the final mixed solution was used immediately.

The electrospinning setup contained a syringe pump (New Era Pump Systems, Inc., Farmingdale, NY, USA), a power supply (DC voltage source, Gamma High Voltage Research, Ormond Beach, FL 32174, USA), a moving stage, and a rotating drum collector. The polymeric solutions were dispensed via a 25-gauge needle at a constant flow rate of 0.450 mL/hr. A driving voltage of 7 kV resulted in a stable jet and the non-woven fiber meshes were collected over a rotating drum covered with nonstick aluminum foil, at a tip-to-ground distance of 7 cm and a drum rotating speed of 80 rpm.

### High-Resolution Scanning Electron Microscopy (HRSEM)

High-resolution scanning electron microscopy (Zeiss GeminiSEM 300, Zeiss, Jena, Germany) was performed in a high vacuum mode, at a work distance (WD) of ∼3-4 mm and voltage of 2-3 kV. After electrospinning, each sample was randomly cut at different locations on the electrospun scaffold and coated with Cr using a sputter coater. HRSEM images were analyzed using ImageJ image processing software to determine morphological appearance and average diameter (averaged over *n* = 80 fibers in each sample).

### Hydrophilicity Measurement

The hydrophilicity of the PCL and PCL−Peptide nanofibers was evaluated through water contact angle measurements using a Drop Shape Analyzer (DSA25, KRUSS) optical tensiometer. Briefly, a drop of water was positioned on the PCL and PCL−Peptide nanofibers (2 cm × 2 cm) film. A side-view photograph of the water droplet was captured at 0 s, 1, 2, 3, and 4 min, and the contact angle between the liquid-air and liquid-mesh interfaces was measured.

### Mechanical Testing

For mechanical tests of the fabricated scaffolds, a universal testing machine (Instron 68SC-2, USA) with a 10 kN load cell capacity was used to determine the tensile strengths and elastic moduli of all scaffolds. Five samples were tested for each scaffold (n = 5). All samples were cut into rectangular films 20 mm in length and 10 mm in width. The thickness of the samples was measured using a film thickness gauge. The samples were gripped on both sides by adhesive tape and their gauge length was 10 mm each. All samples were pulled using a crosshead speed of 5 mm/min.

### Primary cell culture – Keratinocytes and Fibroblasts

Ethical approval for obtaining skin biopsies, and consent forms were obtained and approved by our institution (IRB 4318-17-SMC). A split-thickness biopsy, approximately 2 cm², was obtained from the patient’s non-burned area via surgical procedure. Biopsies were rinsed in PBS supplemented with Penicillin-Streptomycin-Amphotericin B solution for 15 minutes. Keratinocytes were isolated from the biopsy by chopping and incubating the biopsy in trypsin/ethylenediaminetetraacetic acid solution at 37 °C. Keratinocytes were seeded on irradiated 3T3-J2 feeder cells and cultured in keratinocyte growth medium as previously described.^[66]^ Fibroblasts were isolated from the biopsy by attaching the biopsy to a culture plate with the basal membrane facing down for 30 min at 37 °C, then supplemented with DMEM and 10% BCS.

### CEA Sheets

Keratinocytes were cultured in a six-well plate on an irradiated feeder cell layer (i3T3-J2) until confluence. CEA sheets were produced by detaching the cultured keratinocytes using Dispase II and mounting them on fatty gauze backings.

### Cell Viability Measurements

Mixed human skin cells, extracted from a skin biopsy, keratinocytes (1E+05) and fibroblasts (5E+04), were seeded on 1 cm² PCL, PCL-2.5%Fmoc-FRGD, and PCL-20%Fmoc-FRGD electrospun fibers. Cells were cultured with KGM in a 24-well cell-repellent plate. Cell viability was determined by the XTT (2,3-bis-(2-methoxy-4-nitro-5-sulfophenyl)-2H-tetrazolium-5-carboxanilide) test as specified by the supplier (Sartorius) on days 1, 4, 8, and 11 in quadruplicates, with a set of blanks included. Briefly, the activation solution was mixed with XTT reagent (1:500). The reaction solution was then added to each well and incubated for 2 hours at 37°C, 10% CO_2_. Absorbance was determined at 450 nm and 690 nm; net absorbance (A450–A690) was calculated, and blank values were subtracted.

### Scaffolds cross-section and staining

Mixed human skin cells, extracted from a skin biopsy, keratinocytes (2.25E+05) and fibroblasts (1.125E+05), were cultured on 1 cm² PCL and PCL-2.5% Fmoc-FRGD electrospun fibers. At the end of the *iv-vivo* experiment, samples were fixed in 7.5% paraformaldehyde and 8-μm thick cross-sections were prepared. Cells morphology was examined using hematoxylin and eosin (H&E) staining, while cytokeratin (CK10) was used for immunohistochemical staining. Analysis was conducted with Olympus VS200 Slide Scanner (Tokyo, Japan).

### In-vivo Wound Healing Examination

Mouse experiments were conducted under our institution’s approval (SMC-IL-2405-112-4). Mixed human skin cells, extracted from a skin biopsy - keratinocytes (2.25E+05) and fibroblasts (1.125E+05) - were seeded on 1 cm² PCL and PCL-2.5% Fmoc-FRGD scaffolds and grown for 10 days. Keratinocytes were also grown on a feeder layer to produce CEA sheets. A full-thickness skin excision (1 cm²) was made in the center of the back of Nude mouse (male, 9 weeks old) (n=4). Four groups were included: (a) No treatment, control. (b) PCL + skin cells. (c) PCL-2.5% Fmoc-FRGD + skin cells. (d) CEA sheet. The wound healing process was evaluated by visual inspection and wound measurements. Each wound was photographed with a ruler present, and the wound size was measured using ImageJ software on days 0,4,8,11,15 and 19 after graft placement. The wound area of each mouse was normalized to the baseline measurement on day 0. On day 20, the mouse was sacrificed, and a skin excision was made from the closed wound area for histological analysis of the healed skin. The healed skin tissues were further evaluated using H&E histological staining and cytokeratin (CK10) immunohistochemical staining.

### Statistical Analysis

All data analyses were performed using Prism 9.5.1 (GraphPad) and expressed as the mean ± SD based on at least three independent experiments. Specific comparisons are indicated in the respective figure legends.

## Supporting information

Supplementary Information - Cohen-Gerassi et al

## Acknowledgments

This work was supported by the European Research Council (ERC), under the European Union’s Horizon 2020 research and innovation program (grant agreement no. 948 102) (L.A.-A.), the Israel Science Foundation (ISF grant: 2422/24) (L. A.-A.), and the Zimin foundation. D.C.-G. acknowledges the Marian Gertner Institute for Medical Nano Systems at Tel Aviv University. D.C.-G. gratefully acknowledges the support of the Colton Foundation. The authors acknowledge the Jan Koum Center for Nanoscale Systems of Tel Aviv University for the use of instruments and staff assistance. We thank the members of the Adler-Abramovich, Di Segni, and Sitt groups for the help

## References

[1] M. P. Rowan, L. C. Cancio, E. A. Elster, D. M. Burmeister, L. F. Rose, S. Natesan, R. K. Chan, R. J. Christy, K. K. Chung, Critical Care 2015 19:1 2015, 19, 1.

[2] Z. Hussain, H. E. Thu, M. Rawas-Qalaji, M. Naseem, S. Khan, M. Sohail, J Drug Deliv Sci Technol 2022, 68, 103092.

[3] C. Cuono, R. Langdon, J. McGuire, Lancet 1986, 1, 1123.

[4] I. Brockmann, J. Ehrenpfordt, T. Sturmheit, M. Brandenburger, C. Kruse, M. Zille, D. Rose, J. Boltze, Stem Cells Int 2018, 2018.

[5] T. Leclerc, C. Thepenier, P. Jault, E. Bey, J. Peltzer, M. Trouillas, P. Duhamel, L. Bargues, M. Prat, M. Bonderriter, L. Jean-Jacques, Cell Prolif 2011, 44, 48.

[6] B. Homsombath, R. F. Mullins, C. Brandigi, Z. Hassan, S. Fagan, B. Craft-Coffman, T. Olaveson, P. Fidler, C. Cramer, J. Hershman, Journal of Burn Care & Research 2023, 44, 170.

[7] M. H. Desai, J. M. Mlakar, R. L. McCauley, K. M. Abdullah, R. L. Rutan, J. Paul Waymack, M. C. Robson, D. N. Herndon, J Burn Care Rehabil 1991, 12, 540.

[8] M. Faure, G. Mauduit, D. Schmitt, J. Kanitakis, A. Demidem, J. Thivolet, British Journal of Dermatology 1987, 116, 161.

[9] A. A. Meyer, A. Manktelow, M. Johnson, S. Deserres, S. Herzog, H. D. Peterson, J Trauma 1988, 28, 1054.

[10] B. A. Cairns, S. DeSerres, L. A. Brady, C. S. Hultman, A. A. Meyer, M. R. Madden, B. A. Pruitt, K. J. Farrell, R. G. Tompkins, J Trauma 1995, 39, 75.

[11] F. Zhang, C. Hu, Q. Kong, R. Luo, Y. Wang, ACS Appl Mater Interfaces 2019, 11, 37147.

[12] T. Guan, J. Li, C. Chen, Y. Liu, Advanced Science 2022, 9, 2104165.

[13] M. Aviv, D. Cohen-Gerassi, A. A. Orr, R. Misra, Z. A. Arnon, L. J. W. Shimon, Y. Shacham-Diamand, P. Tamamis, L. Adler-Abramovich, Int J Mol Sci 2021, 22, 9634.

[14] M. Halperin-Sternfeld, M. Ghosh, R. Sevostianov, I. Grigoriants, L. Adler-Abramovich, Chemical Communications 2017, 53, 9586.

[15] P. Chakraborty, M. Aviv, F. Netti, D. Cohen-Gerassi, L. Adler-Abramovich, Macromol Biosci 2022, 22, 2100439.

[16] M. Aviv, M. Halperin-Sternfeld, I. Grigoriants, L. Buzhansky, I. Mironi-Harpaz, D. Seliktar, S. Einav, Z. Nevo, L. Adler-Abramovich, ACS Appl Mater Interfaces 2018, 10, 41883.

[17] D. Cohen-Gerassi, Z. A. Arnon, T. Guterman, A. Levin, M. Ghosh, M. Aviv, D. Levy, T. P. J. Knowles, Y. Shacham-Diamand, L. Adler-Abramovich, Chemistry of Materials 2020, 32, 8342.

[18] L. Schnaider, M. Ghosh, D. Bychenko, I. Grigoriants, S. Ya’ari, T. Shalev Antsel, S. Matalon, R. Sarig, T. Brosh, R. Pilo, E. Gazit, L. Adler-Abramovich, ACS Appl Mater Interfaces 2019, 11, 21334.

[19] G. Fichman, C. Andrews, N. L. Patel, J. P. Schneider, Advanced Materials 2021, 33, 2103677.

[20] B. Jeschke, J. Meyer, A. Jonczyk, H. Kessler, P. Adamietz, N. M. Meenen, M. Kantlehner, C. Goepfert, B. Nies, Biomaterials 2002, 23, 3455.

[21] M. Alipour, M. Baneshi, S. Hosseinkhani, R. Mahmoudi, A. Jabari Arabzadeh, M. Akrami, J. Mehrzad, H. Bardania, J Biomed Mater Res A 2020, 108, 839.

[22] E. Ruoslahti, Annu Rev Cell Dev Biol 1996, 12, 697.

[23] J. Gailit, R. A. F. Clark, Curr Opin Cell Biol 1994, 6, 717.

[24] T. Kai, L. Aviad, G. Lihi, Adler-Abramovichab Ehud, Chem. Soc. Rev. 2016, 45, 3935.

[25] D. Cohen-Gerassi, O. Messer, G. Finkelstein-Zuta, M. Aviv, B. Favelukis, Y. Shacham-Diamand, M. Sokol, L. Adler-Abramovich, Adv Healthc Mater 2024.

[26] T. N. Tikhonova, V. S. Kolmogorov, R. V. Timoshenko, A. N. Vaneev, D. Cohen-Gerassi, L. A. Osminkina, P. V. Gorelkin, A. S. Erofeev, N. N. Sysoev, L. Adler-Abramovich, E. A. Shirshin, Cells 2022, 11, 4137.

[27] A. Mahler, M. Reches, M. Rechter, S. Cohen, E. Gazit, Advanced Materials 2006, 18, 1365.

[28] V. Jayawarna, M. Ali, T. A. Jowitt, A. F. Miller, A. Saiani, J. E. Gough, R. V. Ulijn, Advanced Materials 2006, 18, 611.

[29] T. N. Tikhonova, N. N. Rovnyagina, Z. A. Arnon, B. P. Yakimov, Y. M. Efremov, D. Cohen-Gerassi, M. Halperin-Sternfeld, N. V. Kosheleva, V. P. Drachev, A. A. Svistunov, P. S. Timashev, L. Adler-Abramovich, E. A. Shirshin, Angewandte Chemie International Edition 2021, 60, 25339.

[30] R. Orbach, L. Adler-Abramovich, S. Zigerson, I. Mironi-Harpaz, D. Seliktar, E. Gazit, Biomacromolecules 2009, 10, 2646.

[31] H. W. Ju, O. J. Lee, J. M. Lee, B. M. Moon, H. J. Park, Y. R. Park, M. C. Lee, S. H. Kim, J. R. Chao, C. S. Ki, C. H. Park, Int J Biol Macromol 2016, 85.

[32] S. Chen, B. Liu, M. A. Carlson, A. F. Gombart, D. A. Reilly, J. Xie, Recent advances in electrospun nanofibers for wound healing, Vol. 12, 2017.

[33] S. Zhang, W. Yang, W. Gong, Y. Lu, D. G. Yu, P. Liu, RSC Adv 2024, 14, 14374.

[34] R. Xu, Y. Fang, Z. Zhang, Y. Cao, Y. Yan, L. Gan, J. Xu, G. Zhou, Materials 2023, Vol. 16, Page 5459 2023, 16, 5459.

[35] S. G. Kumbar, S. P. Nukavarapu, R. James, L. S. Nair, C. T. Laurencin, Biomaterials 2008, 29, 4100.

[36] F. Hajiali, S. Tajbakhsh, A. Shojaei, Polymer Reviews 2018, 58, 164.

[37] P. Mulinti, J. E. Brooks, B. Lervick, J. E. Pullan, A. E. Brooks, Hemocompatibility of Biomaterials for Clinical Applications: Blood-Biomaterials Interactions 2017, 253.

[38] S. Bayat, N. Amiri, E. Pishavar, F. Kalalinia, J. Movaffagh, M. Hahsemi, Life Sci 2019, 229.

[39] J. Hermosilla, E. Pastene-Navarrete, F. Acevedo, Pharmaceutics 2021, 13.

[40] M. Gizaw, J. Thompson, A. Faglie, S. Y. Lee, P. Neuenschwander, S. F. Chou, Bioengineering 2018, 5, 9.

[41] Y. Dror, J. Kuhn, R. Avrahami, E. Zussman, Macromolecules 2008, 41.

[42] S. Parham, A. Z. Kharazi, H. R. Bakhsheshi-Rad, H. Ghayour, A. F. Ismail, H. Nur, F. Berto, Materials 2020, 13, 2153.

[43] D. Rachmiel, I. Anconina, S. Rudnick-Glick, M. Halperin-Sternfeld, L. Adler-Abramovich, A. Sitt, Int J Mol Sci 2021, 22, 2425.

[44] A. Tambralli, B. Blakeney, J. Anderson, M. Kushwaha, A. Andukuri, D. Dean, H. W. Jun, Biofabrication 2009, 1.

[45] Q. L. Loh, C. Choong, Tissue Eng Part B Rev 2013, 19, 485.

[46] R. Orbach, I. Mironi-Harpaz, L. Adler-Abramovich, E. Mossou, E. P. Mitchell, V. T. Forsyth, E. Gazit, D. Seliktar, Langmuir 2012, 28, 2015.

[47] A. Barth, Biochimica et Biophysica Acta (BBA) - Bioenergetics 2007, 1767, 1073.

[48] R. Xu, H. Xia, W. He, Z. Li, J. Zhao, B. Liu, Y. Wang, Q. Lei, Y. Kong, Y. Bai, Z. Yao, R. Yan, H. Li, R. Zhan, S. Yang, G. Luo, J. Wu, Sci Rep 2016, 6.

[49] C. C. Wu, C. K. Wei, C. C. Ho, S. J. Ding, Materials 2015, 8, 684.

[50] D. J. Rubin, H. T. Nia, T. Desire, P. Q. Nguyen, M. Gevelber, C. Ortiz, N. S. Joshi, Biomacromolecules 2013, 14, 3370.

[51] M. S. Liberato, S. Kogikoski, E. R. Da Silva, D. R. De Araujo, S. Guha, W. A. Alves, J Mater Chem B 2016, 4, 1405.

[52] J. F. M. Manschot, A. J. M. Brakkee, J Biomech 1986, 19, 511.

[53] N. Arnold, J. Scott, T. R. Bush, J Tissue Viability 2023, 32, 286.

[54] T. T. Tran, Z. A. Hamid, K. Y. Cheong, J Phys Conf Ser 2018, 1082.

[55] J. Candiello, M. Balasubramani, E. M. Schreiber, G. J. Cole, U. Mayer, W. Halfter, H. Lin, FEBS Journal 2007, 274, 2897.

[56] J. G. Meinhart, J. C. Schense, H. Schima, M. Gorlitzer, J. A. Hubbell, M. Deutsch, P. Zilla, Tissue Eng 2005, 11.

[57] J. C. Schense, J. A. Hubbell, Journal of Biological Chemistry 2000, 275.

[58] E. Ruoslahti, Annu Rev Cell Dev Biol 1996, 12, 697.

[59] A. A. Khalili, M. R. Ahmad, Int J Mol Sci 2015, 16, 18149.

[60] N. Maas-Szabowski, A. Shimotoyodome, N. E. Fusenig, J Cell Sci 1999, 112, 1843.

[61] Y. Wang, G. Wang, X. Luo, J. Qiu, C. Tang, Burns 2012, 38, 414.

[62] H. Zarkoob, S. Bodduluri, S. V Ponnaluri, J. C. Selby, E. A. Sander, E. A. Sander,; J C Selby, Cell Mol Bioeng 2015, 8, 32.

[63] J. Li, J. N. Chen, Z. X. Peng, N. B. Chen, C. B. Liu, P. Zhang, X. Zhang, G. Q. Chen, ACS Applied Materials and Interfaces 2023, 15, 364.

[64] C. B. Croft, D. Tarin, J Anat 1970.

[65] P. Martin, Science (1979) 1997, 276, 75.

[66] A. Burd, Transplantation 2000, 70, 1551.

