## Supplementary Information - Cohen-Gerassi et al for "Stable, Easy-to-Handle, Fully Autologous Electrospun Polymer-Peptide Skin Equivalent for Severe Burn Injuries"

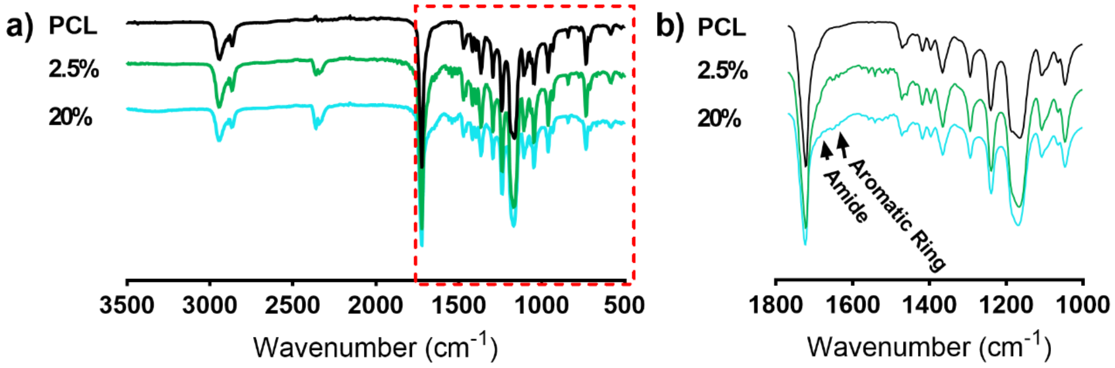


Figure S1. FTIR spectra of the PCL and PCL−peptide nanofibrous matrix.

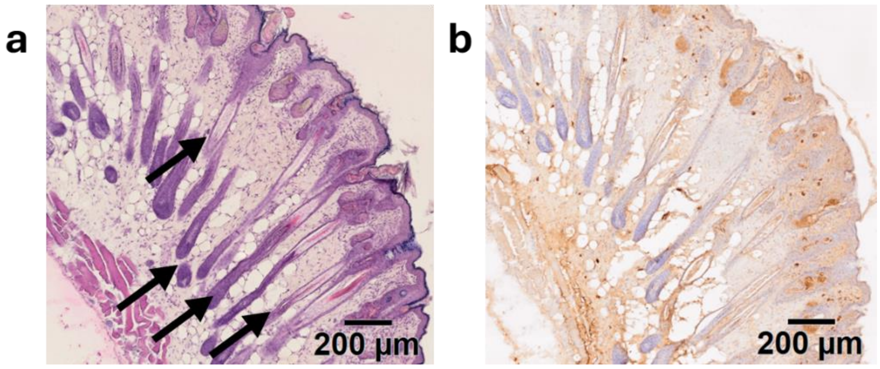


Figure S2. Histological evaluation of healthy skin. (a) H&E staining healthy skin tissues. Black arrows indicate hair follicles (b) Immunohistochemistry CK10 staining of healthy skin tissues.
